# Extracellular matrix remodeling supports *Hydra vulgaris* head regeneration and stem cell invasion

**DOI:** 10.1101/2025.11.07.687297

**Authors:** Ben D. Cox, Jasmine Mah, Angel Perez, Celina E. Juliano

## Abstract

The small freshwater cnidarian *Hydra vulgaris* is a classic model for investigating the genetic regulation of whole-body regeneration, but the underlying cell biology is comparatively underexplored. *Hydra* has a simple body plan consisting of two epithelial monolayers separated by an extracellular matrix (ECM). This ECM contains conserved components such as collagen and laminin, making *Hydra* well suited for dissecting ECM function during regeneration. Following head amputation and wound closure, we observe a retraction of ECM proteins from the wound site, creating a region of low ECM protein accumulation that persists for several days during head regeneration. Several matrix metalloproteinase (MMP) genes are expressed during this process, and MMP inhibition reduces the size of the ECM gap and results in a regenerative outcome with gross morphological defects. We further find that interstitial stem cells (ISCs), which originate in the ectoderm, localize in the regenerating head endoderm near regions of reduced ECM. This suggests that the ECM gap facilitates stem cell invasion to populate the new head with neurons and gland cells. However, inhibition of collagen cross-linking reveals that collagen synthesis is also required for regeneration, indicating that *Hydra* must balance ECM degradation and synthesis to complete regeneration. Together, these findings highlight ECM remodeling as a critical and conserved feature of regeneration.

**Summary statement:** This study uses *Hydra vulgaris*, a highly regenerative freshwater cnidarian, to study remodeling of extracellular matrix proteins and stem cell invasion during tissue regeneration.

## Introduction

Regeneration, defined as the restoration of tissue lost to injury, requires the precise coordination of cell behaviors across time and space. The evolution of the extracellular matrix (ECM) played a key role in the emergence of multicellularity and is integral to coordinating such multicellular and tissue-level behaviors (Brunet and King, 2017; Zhang et al., 2007). Once thought to be a passive scaffold, the ECM is now recognized as a dynamic structure composed of proteins and complex sugars that regulate molecular signaling, morphogenesis, cell migration, and disease (Rozario and DeSimone, 2010). ECM proteins make up a substantial portion of animal tissues, and all metazoans have an ECM based around collagen, the most abundant protein in the animal kingdom (Rozario and DeSimone, 2010; Zhang et al., 2007). Thus, it follows that the ECM should play a key role in regeneration, a process that requires extensive choreography of cell behaviors.

The overall composition of ECM, modulated through secretion and degradation of ECM proteins, must be carefully balanced during embryonic development, homeostasis, wound repair, and regeneration (Walker et al., 2018). Examples from development, such as branching morphogenesis, demonstrate that the function of any ECM component can shift with context, promoting or inhibiting developmental processes (Rozario and DeSimone, 2010). While ECM primarily undergoes fine tuning of its composition during development, complete degradation can occur, such as in the formation of the vertebrate mouth (Chen et al., 2017). Localized removal of the ECM through enzymatic degradation or physical displacement also occurs during development to facilitate cell invasion that is necessary to build animal body plans (Hagedorn et al., 2013; Hiramatsu et al., 2013; Kelley et al., 2014; Nakaya et al., 2008). ECM remodeling plays a prominent role across metazoan regeneration, including whole body regeneration in planarians, amphibian limb regeneration, mammalian digit tip and skin regeneration, and the regeneration of multiple zebrafish structures, including fin, spinal cord, and heart (Calve et al., 2010; Chan et al., 2021; Chen et al., 2015; Constanty et al., 2025; Seifert et al., 2012; Simkin et al., 2015; Tsata et al., 2021). Loss-of-function experiments have shown regeneration-specific expression of diverse collagen subtypes is essential for regeneration in planaria, zebrafish fin and spinal cord, and mammalian skin (Chan et al., 2021; Senk and Djonov, 2021; Volk et al., 2011; Wehner et al., 2017).

Changes in collagen and other ECM components during regeneration define a “transitional matrix” that promotes cell proliferation and differentiation and reduces fibrotic scarring, enabling tissue restoration (Calve et al., 2010; Garcia-Puig et al., 2019; Seifert et al., 2012; Simkin et al., 2015; Tsata et al., 2021). Further changes to the ECM during regeneration include increased expression of ECM-modifying enzymes and glycoproteins including tenascin and fibronectin (Constanty et al., 2025; Garcia-Puig et al., 2019; Tsata et al., 2021). Consistent with the complex, context-dependent requirements for ECM regulation during morphogenesis, regeneration fails in vertebrate tissues with limited regenerative potential when fibrotic scars are not cleared or when ECM proteins such as collagens accumulate excessively (Dent, 1962; Poss et al., 2002; Satoh et al., 2005; Tsata et al., 2021). Despite these important insights, the impact of regeneration-induced ECM dynamics on specific cell behaviors during regeneration, such as the potential role in enabling cell invasion, remain comparatively understudied.

Localized changes in ECM composition are mediated by a variety of remodeling enzymes, the best characterized being the matrix metalloproteinases (MMPs), a large family of genes encoding zinc-dependent proteolytic enzymes. MMPs play key roles in degrading ECM proteins, with most family members possessing at least some collagenase activity, and their activity has been linked to cancer cell invasiveness (Zitka et al., 2010). MMPs are required for regeneration across diverse animal models, underscoring the conserved importance of ECM remodeling in this process, but whether their role in regeneration includes regulating migration and invasion of progenitor cells remains comparatively understudied (Bai et al., 2005; Isolani et al., 2013; LeBert et al., 2015; Simkin et al., 2024; Vinarsky et al., 2005; Yang et al., 1999; Yang and Bryant, 1994).

The small freshwater cnidarian *Hydra vulgaris*, which is capable of whole-body regeneration, has a simple body plan consisting of two epithelial monolayers separated by an ECM (Figure 1A-B). This simplicity makes an excellent system for studying the ECM: with only two epithelial layers, *Hydra* possesses a single continuous ECM, historically called mesoglea, that separates the epithelia and encompasses the animal (Figure 1C-E). *Hydra* ECM contains a layer of fibrillar Collagen I between two layers of non-fibrillar Collagen IV and Laminin, which are in contact with the epithelia (Shimizu et al., 2008) (Figure 1C, D). As the sister group to bilaterians, cnidarians such as *Hydra* occupy a key phylogenetic position, where comparisons with more commonly studied bilaterian organisms can shed light on conserved mechanisms and molecules (Genikhovich and Technau, 2017). Core components of basement membranes and other ECM structures are conserved throughout metazoa: the layers of Collagen IV and Laminin and single layer of Collagen I are similar in organization to the basal laminae and interstitial matrix, respectively, of vertebrates (Özbek et al., 2010; Sarras, 2012).

**Figure 1.**
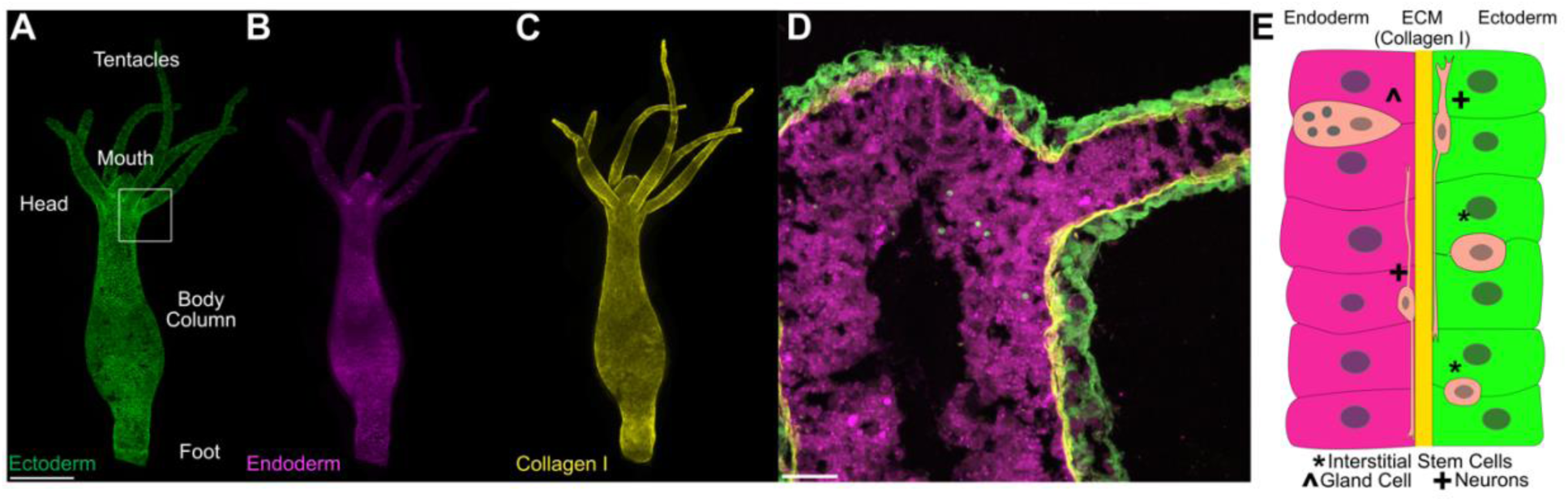
*Hydra vulgaris* as a model for studying extracellular matrix function during regeneration. (A-B) Whole-mount image of a transgenic *Hydra* Tg(*actin1*:GFP)^rs1-ec^;(*actin1*:DsRed)^rs1-en^ (also known as “Watermelon”) expressing GFP (green) in the ectodermal epithelium (A) and RFP (magenta) in the endodermal epithelium (B). The *Hydra* body plan consists of an oral end bearing the mouth and tentacles, an aboral end bearing the foot, and a body column connecting the two poles. The boxed region in A indicates the approximate location of the cryosection shown in D. Scale bar = 500 µm. (C) Image of the same *Hydra* stained with a monoclonal antibody raised against Collagen I (Sarras et al., 1993). Collagen I forms a continuous layer of fibrils along the animal’s entire body column. (D) Cryosection of a Watermelon strain *Hydra* stained with Collagen I antibody, demonstrating that Collagen I forms a distinct layer as part of the ECM that separates the two epithelial cell layers. Scale bar = 100 µm. (E) Schematic illustrating *Hydra’s* two epithelial layers separated by ECM. Interstitial lineage cells are embedded between epithelial cells in both lineages, but interstitial stem cells are restricted to the ectoderm under homeostatic conditions. The interstitial lineage also includes germ cells and nematocytes (not shown).

In addition to the ectodermal and endodermal cell lineages that form the two epithelial layers, *Hydra* contains a third lineage, the interstitial lineage, whose stem cells (interstitial stem cells, or ISCs) are located exclusively in the ectoderm of the body column (Figure 1E). ISCs give rise to three somatic cell types—neurons, gland cells, and nematocytes—as well as germ cells (David, 2012). Although ISCs are restricted to the ectoderm, both neurons and gland cells are present in the endoderm, indicating that ISCs must invade the endoderm to generate these cell types. Radiolabeling experiments support this idea: labeled ectodermal ISCs were detected in the endoderm of uninjured animals, although direct visualization of ISC invasion has not been achieved (Bode et al., 1987). During head regeneration, a large number of head-specific ISC derivates must be replaced (Siebert et al., 2019, 2008). This includes a large population of head-specific gland cells, whose replenishment would require extensive ISC invasion through the ECM into the endoderm, a process that has not been investigated.

Several studies have revealed key roles for the *Hydra* ECM in regeneration. Dissociated cells can normally reaggregate and reform the *Hydra* body plan, but this process is inhibited by antibodies against *Hydra* Collagen I and Laminin (Sarras et al., 1993). While ECM components are known to undergo remodeling during regeneration, and synthesis of new ECM proteins is required for head regeneration, the effect of ECM remodeling on cell behavior, such as epithelial remodeling and ISC invasion, during this process is uncharacterized (Fowler et al., 2000; Lommel et al., 2018; Sarras et al., 1991; Shimizu et al., 2002). In addition, MMPs are required for *Hydra* foot regeneration, but their expression and activity has not been investigated in head regeneration (Leontovich et al., 2000).

Here we examine ECM dynamics during *Hydra* head regeneration to uncover functional roles. We find that after bisection, the regenerating head displays a gap in ECM protein accumulation immediately upon wound closure, which expands over the first two days in an MMP-dependent manner. Pharmacological inhibition of MMP activity disrupts this process and results in gross morphological defects, including failure to pattern new tentacles. Likewise, inhibition of collagen synthesis blocks head regeneration, demonstrating that both ECM degradation and synthesis are required for *Hydra* head regeneration. Finally, during the period of low ECM protein accumulation in the regenerating head, ISCs invade the endoderm at sites of reduced ECM. Together, these findings establish *Hydra* head regeneration as a valuable model for studying ECM dynamics, collective cell migration, and cell invasion during tissue regeneration.

## Results

### Hydra head regeneration proceeds through a phase of reduced ECM accumulation

To investigate ECM dynamics during head regeneration in *Hydra*, we used a previously validated monoclonal antibody against *Hydra* Collagen I to visualize the ECM from 2 hours post-amputation (hpa) through 72 hpa, when head regeneration is complete (Sarras et al., 1993). Fluorescent confocal imaging of whole-mount regenerating *Hydra* revealed a small gap in Collagen I accumulation at the wound site immediately after closure at 2 hpa (Figure 2A) (Shimizu et al., 2002). Although this initial gap had been previously described, its dynamics over the course of head regeneration were unknown. We found that the gap persisted through the first 12 hpa, and then expanded in size between 12 and 36 hpa (Figure 2B-E). Importantly, this ECM gap was not acellular, as nuclei were visible within these regions (Figure 2A). The ECM protein Laminin showed similar restriction from the regenerating head (Figure S1) (Shimizu et al., 2002).

**Figure 2.**
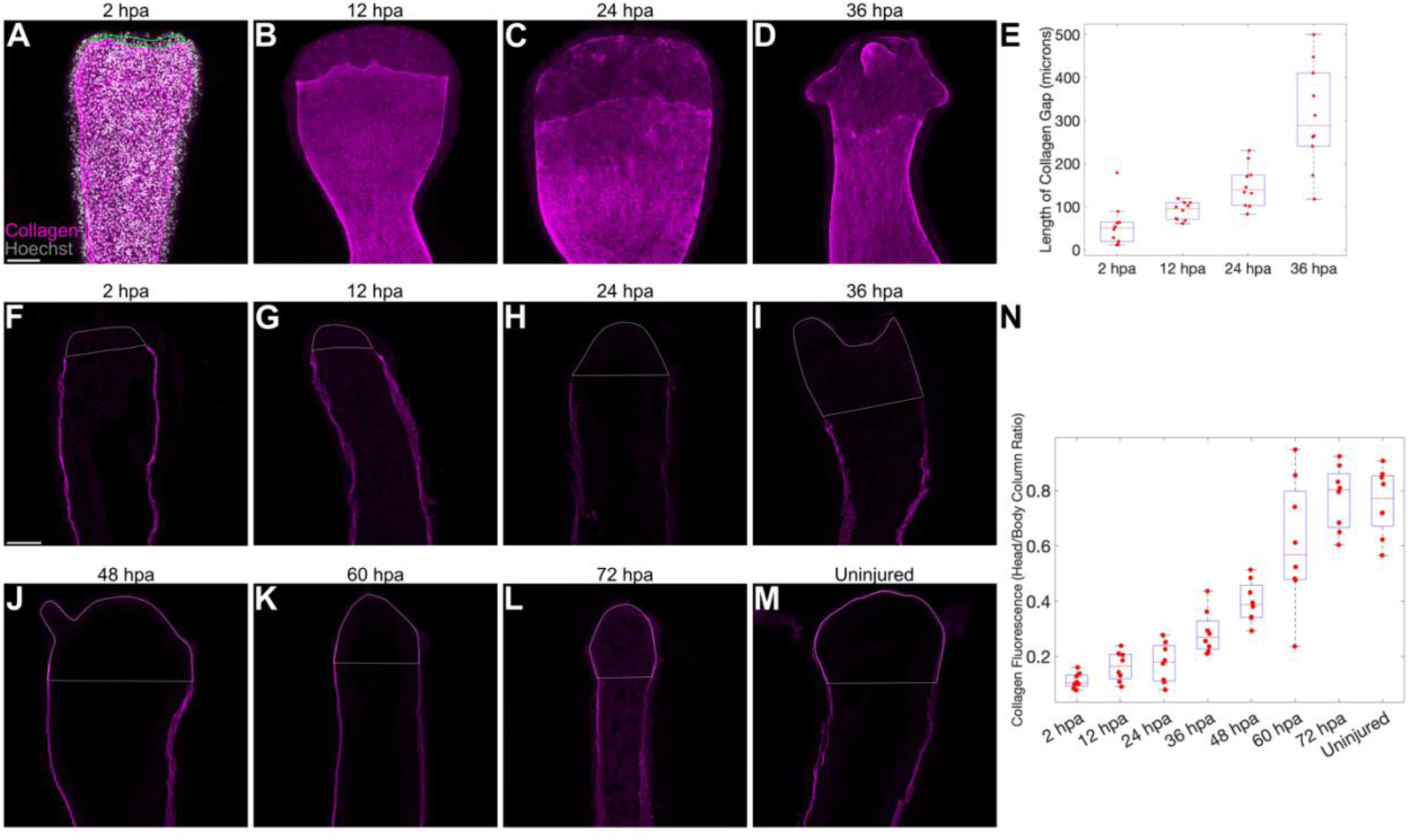
Temporal dynamics of Collagen I loss and renewal after injury. (A) Whole-mount fluorescence imaging shows a Collagen I-depleted region at the site of wound closure and future head regeneration. Hoechst staining (grey) confirms the presence of tissue beyond the oral end of Collagen I staining (marked with green line). Scale bar = 100 µm. (B-D) The Collagen I-depleted region progressively enlarges over the first 36 hpa. (E) Quantification of the average length of the Collagen I gap over the course of head regeneration. (F-M) Time course of head regeneration on cryosectioned animals stained with Collagen I antibody, showing loss and subsequent reappearance of Collagen I. Scale bar = 100 µm. White dashed lines indicate area where head fluorescence intensity was measured and compared to fluorescence intensity in body column ECM. (N) Quantification of the ratio of Collagen I signal intensity in the regenerating head relative to the adjacent body column. Each data point represents the mean of three representative cryosections per animal (n = 8 animals per time point).

After 36 hpa, the ECM gap became increasingly difficult to detect in whole mount images, as the formation of new tentacles increased the thickness of the sample and lifted the head tissue away from the cover slip, reducing the fluorescent signal. To more accurately track Collagen I accumulation in the regenerating head, we imaged representative longitudinal cryosections of regenerating *Hydra* after wound closure and at 12 hour intervals through 72 hpa (Figure 2F-2M). Quantification of Collagen I intensity in the regenerating head relative to the body column showed almost complete exclusion from the wound site immediately after epithelial closure (Figure 2F, 2N). Collagen I head-to-body column ratios remained below half of those in uninjured animals until 48 hpa and then recovered to approximately the level of uninjured animals by 72 hpa (Figure 2N).

### Endodermal MMPs mediate ECM degradation in the regenerating head

We observed that the ECM gap increased in size between 12 and 48 hpa, suggesting active degradation of ECM components during this period. To test if MMP family members, which degrade collagen proteins, contribute to this process, we treated regenerating *Hydra* with the broad-spectrum MMP inhibitor Marimastat and measured the size of the Collagen I-depleted region during head regeneration (Figure 3A-C) (Rasmussen and McCann, 1997). At 24 hpa, the length of the Collagen I-depleted region, measured from the tip of the regenerating head to the border of low Collagen I expression, was significantly reduced (Figure 3C) in animals treated with 50 µM Marimastat (Figure 3B) compared to vehicle controls (0.1% DMSO, Figure 3A). Continuous inhibition through the first 48 hpa resulted in a significant delay in gross morphological regeneration, as measured by the appearance of new tentacle buds (Figure 3D-F).

**Figure 3.**
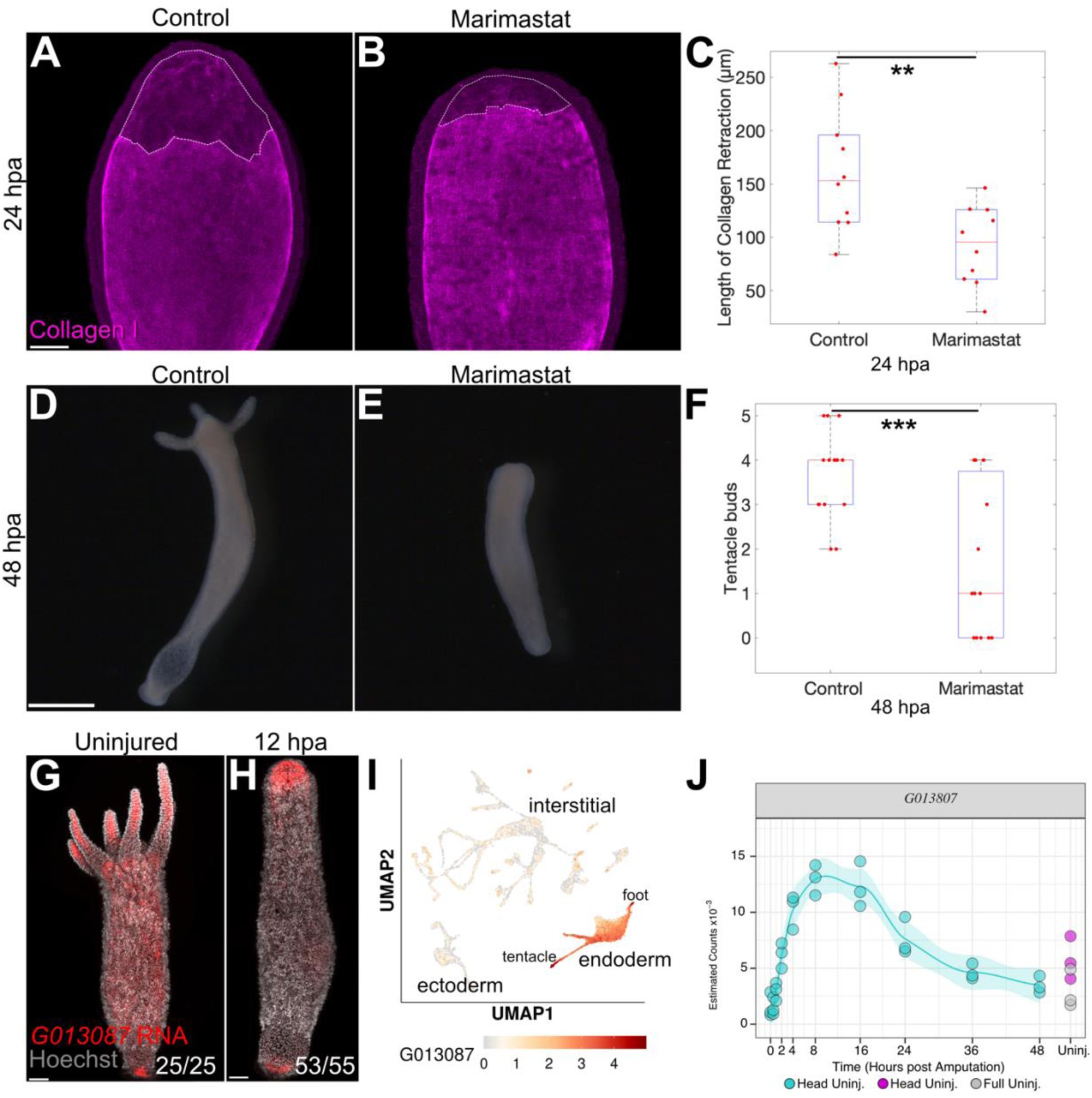
MMP activity is essential for Collagen I degradation at the wound site and for successful morphological regeneration. (A-B) Collagen I antibody staining of 0.1% DMSO-treated (A) and 50 µM Marimastat-treated (B) animals at 24 hpa. Scale bar = 100 µm. White dashed line indicates the measured Collagen I-depleted region. The area of this region was divided by the width of the body column at the aboral end of the gap to determine the average length of retraction. (C) Quantification of the average length of the Collagen I gap. n = 10 animals, p < 0.01 (D-E) Whole-mount images of 0.1% DMSO-treated (D) and 50 µM Marimastat-treated (E) *Hydra* at 48 hpa show significant reduction in tentacle bud formation following MMP inhibition. Scale bar = 500 µm. (F) Quantification of tentacle buds at 48 hpa (D-E). n = 15 animals, p < 0.001. (G– H) Fluorescence in situ hybridization (FISH) for *G013087* in uninjured (G) and regenerating (H) animals. *G013087* is expressed throughout the endoderm in uninjured *Hydra*, with strongest expression in the tentacles and foot, and is upregulated in the regenerating head endoderm by 12 hpa. Scale bars = 100 µm. The ratio in the bottom right of each panel indicates the number of animals showing the depicted expression pattern out of total stained animals. (I) Previously published uninjured animal scRNA-seq UMAP showing *G013087* expression is consistent with the FISH data (Cazet et al., 2023; Siebert et al., 2019). (J) Previously published bulk RNA-seq data shows a significant increase in *G013087* during head regeneration (Wenger et al., 2019).

Because Marimastat broadly inhibits MMP family members, we next sought to identify individual MMPs expressed during head regeneration as candidates for this activity. Previously published transcriptomic data from our lab identified twelve transcripts with UniProt BLAST hits matching members of the MMP family. We searched the sequences of these transcripts for the N-terminal propeptide, catalytic domain, and C-terminal/hemopexin-like domain characteristic of MMP family members (Zitka et al., 2010) (Figure S2. Tables S1-2). Of these transcripts, six had all three domains present (Figure S2A-F), three lacked an N-terminal propeptide (Figure S1G-I), two lacked a hemopexin-like domain (Figure S2J-K), and one lacked all domains characteristic of MMPs (Figure S2L). Because some MMPs lack the hemopexin-like domain and the N-terminal propeptide is removed during activation, we considered the eleven transcripts with a bona fide catalytic domain as candidates for ECM degradation during regeneration (Fig S2A-K).

To assess whether these transcripts could plausibly function in head regeneration, we examined their expression patterns using published transcriptional resources and fluorescence *in situ* hybridization (FISH) (Bonnans et al., 2014). Previously published single cell RNA sequencing (scRNA-seq) data on whole animals and bulk RNA-seq data taken during head regeneration indicated that two transcripts (*G018947*, *G018949*) had very low expression in the uninjured animals and showed no change during regeneration, thus ruling out their function in ECM degradation during head regeneration. For the remaining nine transcripts we observed significant expression in the uninjured and regenerating animal in the published transcriptional data, so we next aimed to test their expression patterns using FISH. One of these candidates (*G004520*) was predicted by scRNA-seq to be broadly expressed across all three cell lineages, but we failed to generate a reliable FISH probe. For the final eight mmp transcripts, we successfully performed FISH on uninjured *Hydra* and animals undergoing head regeneration at 12 hpa (Figure 3G-H, Figure S3). Comparing FISH results of uninjured animals to previously published scRNA-seq data confirmed that localization matched predicted cell type expression (Figure 3G, 3I, Figure S2). Notably, five MMP genes (*G013087*, *G018597*, *G018104*, *G013033*, *G022819*) showed specific expression in the regenerating heads at 12 hpa in at least 75% of the sample (Figure 3H, Figure S3A-E). One of these *mmp* transcripts (*G013087*) had previously been shown to localize to the regenerating head, but the remaining four had not been previously identified as expressed in the regenerating head (Leontovich et al., 2000). All five localized specifically to the endoderm, indicating that ECM degradation is primarily an endoderm-driven process.

We next examined temporal expression dynamics of these MMPs using a previously published bulk RNA-seq time course of head regeneration extending to 48 hpa (Wenger et al., 2019). Of the five transcripts that showed head endoderm localization during regeneration, two (*G013087* and *G018597*) showed significant upregulation during the first 12 hpa (Figure S3A, 2B). The remaining three transcripts showed more modest increases during regeneration, and expression peaked after 12 hpa (Figure S3C-E). Of the remaining three *mmp* transcripts for which we were able to detect expression, two were expressed in gland cells of the interstitial lineage and showed a decrease in expression during regeneration (Figure S3F-G). A third was expressed in both endoderm and interstitial cell types and showed little change in expression during regeneration (Figure S3H). In summary, MMP activity is necessary for the ECM degradation during head regeneration, and its inhibition results in downstream delays in head morphogenesis. Several *mmp* genes show regeneration-specific, head-localized expression in the endoderm, pointing to an endodermal role in driving ECM remodeling during regeneration.

### Endodermal collagen synthesis is required for head regeneration

Our MMP inhibition experiments demonstrate that collagen breakdown is required for *Hydra* head regeneration. Previous studies reported that inhibition of collagen maturation also inhibits head regeneration, but the phenotype was not well characterized (Sarras et al., 1991). To address this gap, we treated *Hydra* with the collagen cross-linking inhibitor dipyridyl for 48 hours and assessed regeneration by counting the appearance of new tentacle buds. While vehicle-treated animals showed no delay in tentacle bud formation, dipyridyl treatment completely inhibited tentacle development and caused animals to adopt a balloon-like morphology (Figure 4A-B, F). After assessing head regeneration at 48 hpa, we released half of the dipyridyl-treated animals from the drug and re-assessed tentacle bud formation at 96 hpa (Figure 4C-F). Approximately half of the released animals recovered and produced tentacle buds similar to untreated animals at 48 hpa, while the remainder failed to regenerate heads, retained the balloon-like phenotype, and became so fragile at the injury site that most ruptured upon fixation and imaging. However, the relatively high rate of 50% recovery observed after drug washout indicates that the regeneration block is likely not due to nonspecific toxicity. To confirm that dipyridyl treatment inhibits collagen maturation in *Hydra*, we performed whole mount immunofluorescence at 24 hpa. Vehicle-treated control animals displayed patchy regions of new collagen synthesis in the regenerating head (figure 4G), whereas dipyridyl-treated animals showed greatly reduced collagen signal intensity (Figure 4H-I).

**Figure 4.**
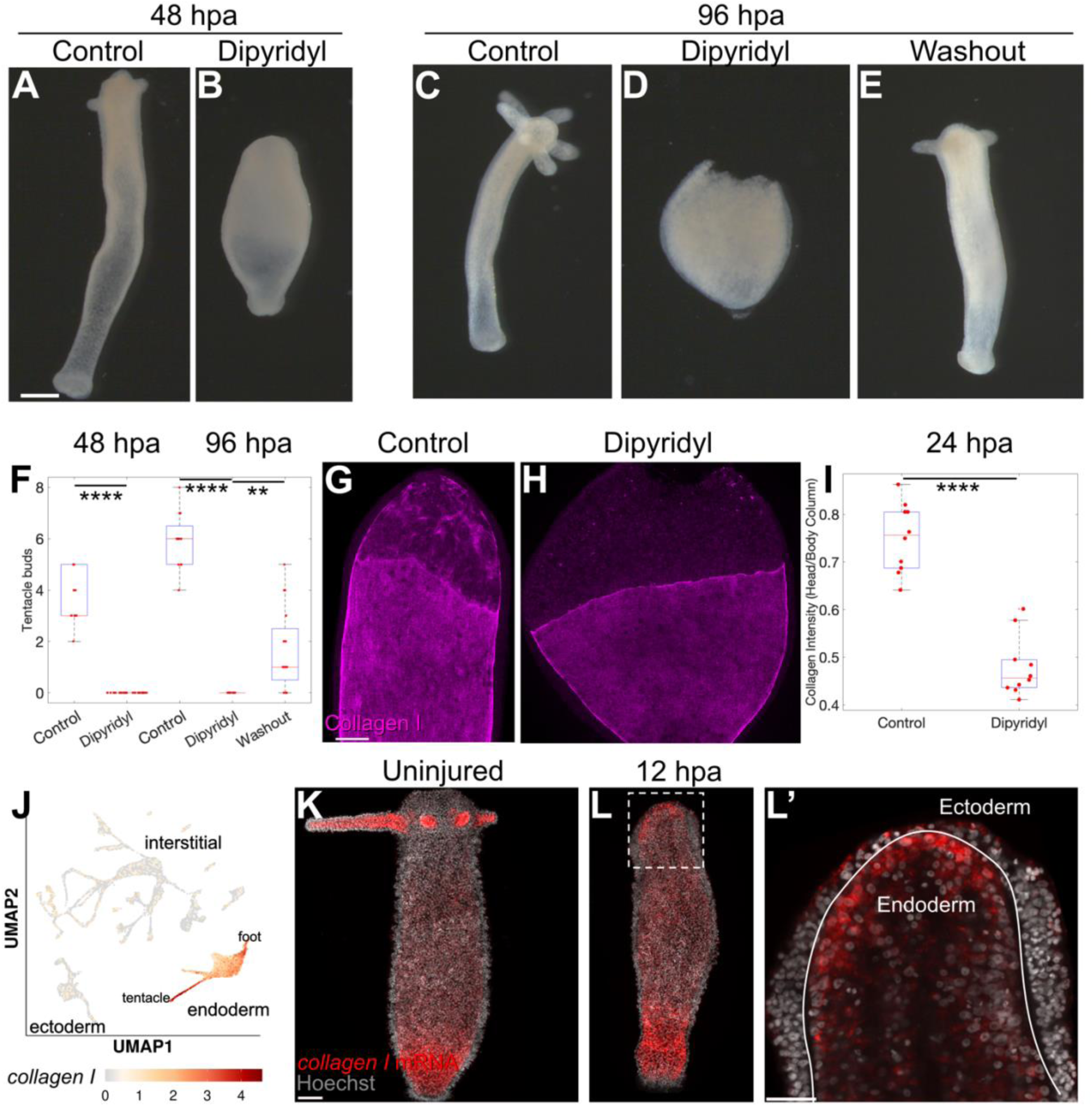
Collagen triple helix formation is essential for *Hydra* head regeneration and new collagen deposition in the regenerating head. (A-B) *Hydra* treated with vehicle control 0.1% ethanol (A) show normal tentacle formation at 48 hpa, while 100 µM dipyridyl-treated animals (B) fail to form tentacle buds and display a stretched, balloon-like morphology of the head and body column. Scale bar = 100 µm. (C-E) At 96 hpa, 0.1% ethanol-treated *Hydra* (C) continue normal tentacle development, while dipyridyl-treated animals (D) still lack tentacles, and the regenerating head tissue becomes fragile and prone to rupture. When animals are switched from dipyridyl to 0.1% ethanol at 48 hpa and allowed to recover for an additional 48 h (E), approximately half form tentacle buds. (F) Quantification of tentacle bud formation at 48 and 96 hpa for animals shown in panels A-E, n = 16 animals (8 each for 96 hpa dipyridyl and washout experiments), **** p < 0.0001, ** p < 0.01. (G-H) Collagen fibrils, visualized by antibody staining, are detectable in the regenerating heads of ethanol treated animals as early as 24 hpa (G), while dipyridyl-treated animals (H) show significantly reduced signal and lack organized fibrillar structure. Scale bar = 100 µm. (I) Quantification of collagen fluorescence intensity, expressed as the ratio of head to body column fluorescence, reveals significantly reduced collagen in dipyridyl treated animals. n = 10 animals, **** p < 0.0001. (J) UMAP of *collagen I* (*G020291*) mRNA expression using data from Siebert et al. 2019 and Cazet et al. 2023, showing highest expression in the endoderm, with enrichment at the foot and tentacle. (K-L) FISH for *collagen I* mRNA in uninjured animals (K) show brightest fluorescence in foot and tentacle endoderm, consistent with UMAP shown in (J), while in 12 hpa animals (L) fluorescence is brightest in the foot and the regenerating head. (L’) Zoom of boxed region in L. Solid white line traces gap in Hoechst nuclear staining where ECM is present. Head endoderm stains strongly, while red fluorescence outside of endoderm is predominantly extracellular and likely reflects probe trapping. Scale bar for K: 200 µm. Scale bar for L’: 50 µm.

We next investigated the source of new collagen synthesis. Earlier studies using colorimetric whole mount *in situ* hybridization suggested that the ectodermal layer expresses *collagen I* transcripts, but our scRNA-seq data showed expression primarily in the endoderm (Figure 4J) (Cazet et al., 2023; Shimizu et al., 2002). To resolve this discrepancy, we performed FISH for *collagen I* (*G020291*) on uninjured and 12 hpa regenerating animals and with this increased resolution confirmed expression in endodermal cells (Figure 4K-L’). Expression was ubiquitous throughout the endoderm, but stronger in the foot and tentacles of uninjured *Hydra*, consistent with the scRNA-seq data. We also observed strong expression in the regenerating *Hydra* head at 12 hpa. Together, these findings indicate that endodermal epithelium is the primary driver of both collagen breakdown, via expression of MMP enzymes, and new collagen synthesis, and that pharmacological inhibition of either process blocks head regeneration.

### Stem cell invasion during regeneration correlates with low levels of collagen

Our pharmacological studies indicate that successful head regeneration requires a precise balance between collagen breakdown and new synthesis. We further hypothesized that ECM degradation facilitates the movement of ISCs from the ectoderm into the endoderm, where they could replenish lost mucous gland cell populations during regeneration (Figure 5A). To test this, we visualized ISCs during regeneration using the transgenic line Tg(*nanos*:eGFP)^tb1-in^, which expresses eGFP under the control of the *nanos* promoter (Hemmrich et al., 2012). As expected, whole mount imaging of eGFP-positive ISCs in the uninjured animal were distributed broadly along the body column but excluded from the foot, head, and tentacles (Figure 5B). Longitudinal cryosections confirm that under homeostatic conditions ISCs are restricted to the ectoderm of the body column and are not observed in the endoderm (Figure 5C-D).

**Figure 5.**
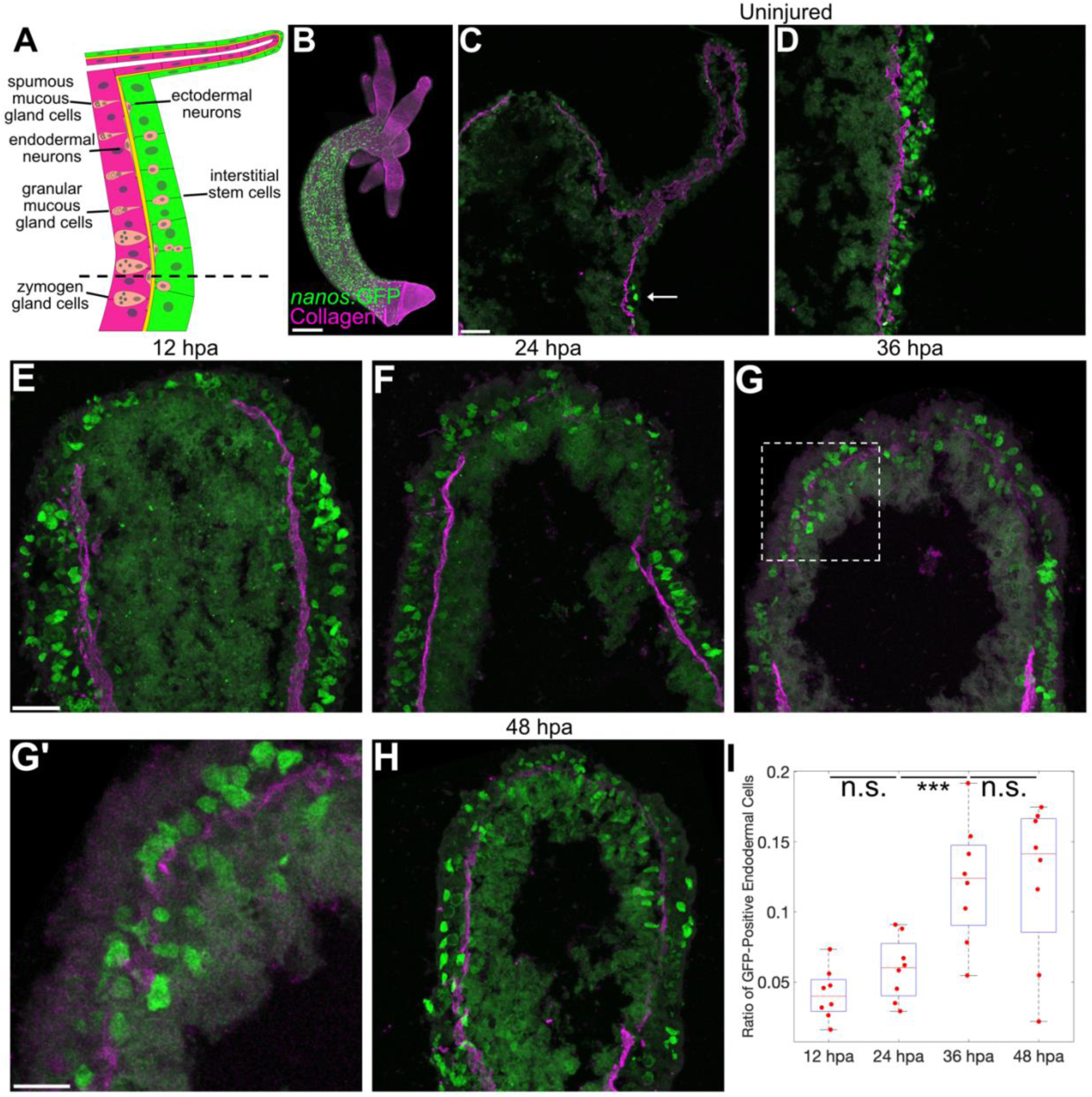
Interstitial Stem Cells (ISCs) invade the endoderm at sites of reduced Collagen I localization during head regeneration. (A) Illustration showing location of ISCs and their derivatives in the oral half of the *Hydra* body column. ISCs are located in the body column ectoderm (green) but excluded from the tentacles, head, and endoderm (magenta). Neurons are located in both the endoderm and ectoderm. Zymogen gland cells are located in the mid-body column endoderm, while spumous and mucous gland cells are located in the upper body column endoderm. Black dashed line indicates typical plane of bisection in injury experiments. (B) Whole mount image of fixed *Hydra* stained with Collagen I antibody (magenta) and expressing eGFP in ISCs under the control of the *nanos* promoter (Tg(*nanos*:eGFP)^tb1-in^). The head, tentacles, and foot lack ISCs. Scale bar = 200 µm. (C) Tissue cryosection of an uninjured *Hydra* shows a lack of eGFP-positive ISCs in the head, tentacles, and endoderm. Arrow: eGFP-positive ISCs located aboral to the head and tentacles. Scale bar for C and D = 50 µm. (D) Tissue cryosection from the body column of an uninjured *Hydra* shows that the ectoderm is dense with eGFP-positive ISCs, while the endoderm lacks ISCs. (E-H) Representative cryosections of eGFP-positive ISC distribution in *Hydra* head regeneration at 12 (E), 24 (F), 36 (G), and 48 (H) hpa. Scale bar for E-G, H = 50 µm. (G’) Zoom of boxed region in G. Presence of eGFP-positive cells in endoderm corresponds to regions of reduced Collagen I antibody staining. Scale bar = 20 µm. (I) Quantification of eGFP-positive endodermal cells in the regenerating head at time points shown in E-H. n = 8 animals, p < 0.001. n.s.: not significant.

To test the hypothesis that low collagen levels facilitate ISC invasion during regeneration, we quantified the presence of eGFP-positive cells in the endoderm within 200 µm of the regenerating head tip at successive 12-hour intervals after injury. Upon bisection, the regenerating head has a large population of ISCs due to body axis realignment, as what was once the middle of the body column is now the presumptive head (Figure 5E). At 12 hpa, very few ISCs were present in the regenerating head endoderm (Figure 5E). However, at successive 12-hour intervals, the percentage of endodermal cells expressing eGFP progressively increased, peaking at 36 hpa (Figure 5F-I). eGFP in the endoderm became more difficult to detect in later time points as eGFP-positive stem cells differentiate, we therefore did not image past 48 hpa.

After 48 hpa, eGFP expression in interstitial derivatives in the endoderm is lost and the homeostatic distribution of ISCs, excluded from the head, is reestablished. Together, these findings show that ISC invasion from the ectoderm into the endoderm coincides with reduced ECM levels in the regenerating head, supporting the hypothesis that the ECM gap facilitates large-scale stem cell invasion during regeneration.

## Discussion

Using fluorescence imaging and pharmacological inhibitors, we have demonstrated that Collagen I in the regenerating *Hydra* head undergoes significant remodeling over the course of 72 hours post-amputation, and that both enzymatic remodeling and new collagen synthesis are essential to regeneration success. Further, we show that ECM remodeling is driven by transcriptional changes in the endodermal epithelial layer. During the period of reduced Collagen I localization in the head, ISCs invade from the ectoderm into the endoderm, then differentiate to replace cell types lost to amputation.

We propose a model for the role of ECM remodeling in *Hydra* head regeneration (Figure 6). In the uninjured head, head-specific gland cells reside in the endoderm (Siebert et al., 2008), and several distinct neuron subtypes are found in both epithelial layers (Little et al., 2025) (Figure 6A). Head amputation removes these head-specific populations and exposes the tissue layers to the environment (Figure 6B). Wound closure rapidly restores epithelial integrity but leaves a small gap in ECM protein localization between the ectodermal and endodermal layers of the head (Figure 6C). Following injury, what was previously body column tissue becomes the new oral region of the animal that will be remodeled to form the regenerating head. ISCs within the ectoderm are therefore positioned to replenish lost cell types by invading the endoderm. During this phase, an initial gap in Collagen I localization at the wound site expands due to endoderm-derived MMP transcription and activity (Figure 6C-D). ISCs invade the endoderm at Collagen I-depleted regions of the regenerating head (Figure 6D-E), where they differentiate to replace endodermal neurons and gland cells. Subsequent synthesis of new Collagen I and other ECM components restores the ECM to its homeostatic state (Figure 6F). Notably, transcriptional upregulation of collagen and other ECM components is concurrent with ECM-degrading enzymes, suggesting that the delay between ECM transcript expression and new ECM protein accumulation in the regenerating head is the result of post-transcriptional and post-translational modifications of ECM mRNA and proteins.

**Figure 6.**
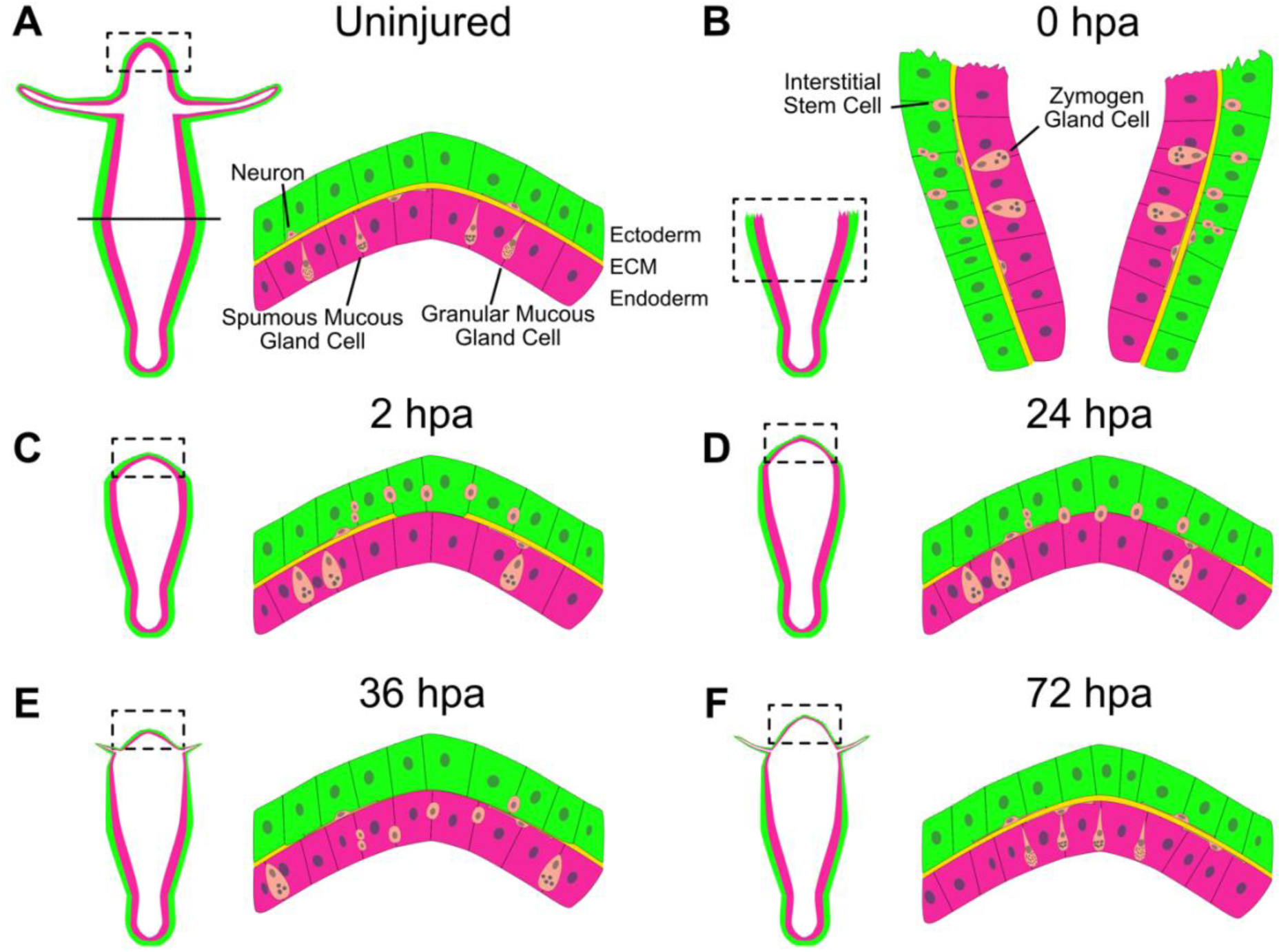
Model for extracellular matrix remodeling and interstitial stem cell invasion during *Hydra* head regeneration. For all panels, right image is the zoomed- in view of the region depicted in the black dashed box at left. (A) *Hydra* has interstitial cell derivatives (peach) embedded within the two epithelial layers. Neurons are present within the ectoderm (green) and endoderm (magenta), while mucous gland cells types are present only in the head endoderm. Solid black line indicates plane of bisection. (B) After oral half amputation, mid-body column tissue will be remodeled into the new head during regeneration. ISCs in the ectoderm are poised to replace lost gland cell and neuron subtypes. (C) After wound closure, a small gap in the ECM (yellow) remains at the site of wound closure. (D) As head regeneration proceeds, the ECM gap grows as a result of MMP activity, and ISCs begin to invade the endoderm. (E) Some ISCs have invaded the endoderm, while others will remain in the ectoderm to replace lost ectodermal interstitial cell types. (F) After head regeneration is complete, new ECM proteins have filled the gap in the regenerating head, and all ISCs in the head have differentiated. Epithelial morphogenesis of the head and later proliferation will push body column-specific cell types, such as zymogen gland cells, away from the mouth.

Collagen remodeling and metalloproteinase activity during *Hydra* regeneration have been described previously (Deutzmann et al., 2000; Leontovich et al., 2000; Shimizu et al., 2002; Yan et al., 1995), and our findings extend these studies in several important ways. We identify a prolonged phase of Collagen I reduction during head regeneration and demonstrate that this period coincides with dynamic MMP expression and activity. Whereas prior work focused on a single MMP (*G13087*) or non-MMP family members, our transcriptomic analysis revealed five MMP-related genes expressed during head regeneration, four of which were previously uncharacterized. Inhibition of MMP activity suppressed the expansion of the ECM-depleted region and led to regeneration defects, confirming a functional role for these enzymes. Finally, by integrating single-cell transcriptomics with FISH, we demonstrate that *collagen I* transcripts are predominantly produced by endodermal epithelial cells, rather than the ectoderm as previously suggested by colorimetric in situ hybridization (Deutzmann et al., 2000).

In other animals, both migrating cells and their surrounding environment can contribute to *mmp* expression and ECM degradation to enable cell invasion during development and regeneration. In *C. elegans*, invading anchor cells express multiple *mmp* family members to promote basement membrane degradation (Kelley et al., 2019). Conversely, during planarian regeneration, unidentified differentiated cells rather than *smedwi-1*-positive migrating stem cells express *mmp*s (Abnave et al., 2017). Finally, zebrafish cardiac regeneration involves *mmp* expression by both invading cardiomyocytes and support cell types, such as fibroblasts and macrophages (Constanty et al., 2025). In *Hydra*, our results indicate that ECM degradation during head regeneration is driven not by expression of *mmp* family members in the invading ISCs themselves, but by *mmp* expression in the endoderm. Co-expression of both *mmp* family members and *collagen I* transcripts in the endoderm suggests that ECM remodeling is an endoderm-driven process that generates an invasion-permissive environment for the ISCs. Although ISC invasion has been inferred during *Hydra* homeostasis via cell labeling experiments, its role in repopulating the regenerating head endoderm had not previously been explored (Bode et al., 1987). We observe ISC accumulation in the regenerating head endoderm coincident with a period of low ECM protein accumulation, suggesting that ECM depletion facilitates large-scale ISC invasion. ECM remodeling-dependent cell invasion has been demonstrated in the regenerating zebrafish heart, suggesting conservation, but this process remains underexplored in most models (Constanty et al., 2025). Our findings establish *Hydra* as a powerful system to study stem cell invasion mechanisms during regeneration.

Given the diverse roles of ECM, it is likely that its remodeling during regeneration serves functions beyond facilitating cell invasion. For example, in addition to acting as a barrier that maintains epithelial integrity, the ECM can function as a signaling center by sequestering morphogens and other molecules that are released by ECM-modifying enzymes (Rozario and DeSimone, 2010). In *Hydra*, overactivation of Wnt signaling leads to reduced ECM elasticity, and Wnt acts upstream of ECM remodeling genes during *Hydra* head regeneration (Veschgini et al., 2023, Campos et al. 2025). Although *Hydra* ECM remodeling is downstream of Wnt signaling, whether the ECM in turn feeds back to influence Wnt or other signaling pathways remains unexplored. Given the rapid loss of ECM integrity upon bisection, it is possible that injury triggers the release of ECM-bound signaling molecules that contribute to initiating regeneration, a possibility that merits future investigation.

Though changes to ECM composition during regeneration have been documented, to the best of our knowledge, the extensive removal of a homeostatic ECM layer that we describe here has not been reported in whole-body regeneration. A comparable case of large-scale ECM depletion does occur in the axolotl limb blastema, which lacks a basement membrane, and where collagen matrix formation marks the end of blastema growth (Rao et al., 2009; Satoh et al., 2011). In mammals, differences in basement membrane composition and thickness correlate with species differences in mouse skin regeneration, where a shift from Collagen I-rich to Collagen III-rich ECM characterizes highly regenerative skin (Gawriluk et al., 2016; Seifert et al., 2012). Collagen I also is considered the main driver of tissue fibrosis across multiple organs (Didangelos et al., 2016). Collagen-containing scars formed early after cardiac injury are eventually degraded during successful zebrafish heart regeneration, whereas failed heart regeneration is associated with retention of Collagen I and likely other collagen subtypes (Bevan et al., 2020). However, in keeping with the many functions individual ECM components play in complex biological processes, Collagen I does not act only as an obstacle to regeneration. Even in models where Collagen I retention contributes to scarring, such as the heart, complete ablation of collagen-expressing cells reduces cardiomyocyte proliferation. This demonstrates that although long-term collagen retention can be detrimental, collagen can still play beneficial roles during earlier stages of regeneration (Sánchez-Iranzo et al., 2018). Here, we demonstrate that Collagen I accumulation is transiently reduced in the regenerating *Hydra* head, suggesting the formation of a transitional matrix with distinct properties from the homeostatic ECM. Such transient remodeling may represent a deeply conserved mechanisms that facilitates regeneration across diverse animals.

Whether a regeneration-specific or transitional ECM exists in *Hydra* after the initial exclusion of Collagen I from the regenerating head, and how it differs from the homeostatic matrix, remains an open question. Work from our lab and others has shown that core ECM components, including laminin and collagen, are transcriptionally upregulated during regeneration (Cazet et al., 2021; Wenger et al., 2019). However, changes at the protein level, where molecules like Collagen I undergo extensive modification to form triple helices, remain largely unexplored. Proteomic studies in *Hydra* have characterized the ECM in uninjured animals and during the early stages of head regeneration, but changes in protein composition after the initial establishment of a head organizer (8-12 hpa) have not yet been documented (Lommel et al., 2018; Petersen et al., 2015). Because the restoration of the homeostatic ECM only occurs at later stages of head regeneration, extending proteomic analyses to these later time points will be critical to identifying features of a potential transitional matrix. Defining the molecular composition of a regenerative ECM in *Hydra* could reveal whether ECM dynamics observed in vertebrate regeneration are conserved across metazoans. Investigating ECM protein dynamics during regeneration will also provide new insights into *Hydra*’s remarkable regenerative capacity and the molecular and evolutionary origin of regeneration.

## Materials and Methods

### Hydra strains and culture

All experiments on non-transgenic *Hydra* were performed on *Hydra vulgaris* AEP strain. The Tg(*nanos*:eGFP)^tb1-in^ strain was generated by Hemmrich and colleagues (Hemmrich et al., 2012). Watermelon strain Tg(actin1:GFP)^rs1-ec^;(actin1:DsRed)^rs1-en^ was generated by Glauber and colleagues (Glauber et al., 2015). All *Hydra* strains were cultured in *Hydra* medium (0.38mM CaCl_2_, 0.32 mM MgSO_4_ X 7H2O, 0.5 mM NaHCO_3_, 0.08 mM K_2_CO_3_) at 18° C, and fed three days per week with freshly hatched *Artemia nauplii* (https://www.brineshrimpdirect.com). All animals were starved for 24 h before experimentation. See Little et al., 2025 for explanation of *Hydra* transgenic line naming scheme.

### Injury

To induce regeneration, *Hydra* were observed under a stereomicroscope, allowed to relax to maximum length, and cut at the midpoint of the body column using a Standard Scalpel Blade #10 (World Precision Instrument 500239).

### Animal fixation

Animals were anesthetized for two minutes in 2% urethane in *Hydra* medium (see *Hydra* strains and culture section), then fixed in 4% paraformaldehyde (16% paraformaldehyde [ThermoFisher 043368.9M] diluted to 1:3 in *Hydra* medium) for 1 h at room temperature or overnight (approximately 16 h) at 4° C.

### Tissue cryosectioning

After fixation, samples were washed 3x for 5-10 minutes per wash in phosphate-buffered saline (PBS) solution, then successively incubated in 10%, 20%, and 30% sucrose solutions in PBS for at least 1 h each. Samples were then arranged in embedding medium (Fisher Tissue Plus OCT Compound 4585) in cryomolds (Sakura TissueTek 4557) and frozen at −70°C. Frozen blocks were sectioned at 25 microns per section (Leica CM1850 UV cryostat) and melted onto slides (Fisher Brand Superfrost Plus 12-550-15). Slides were warmed for 1 h at 37° C prior to staining.

### Antibody staining

Whole mount immunofluorescence: after fixation, animals were washed 3x in PBS for 10 minutes each, then incubated in PBS with 0.5% Triton X-100 2x for 15 minutes each. Blocking was performed for 1 h at room temperature or overnight (approximately 16 hours) at 4° C in a solution of PBS with 0.1% Triton X-100, 10% goat serum, and 1% w/v BSA. Antibody staining was then performed in blocking solution for 1 h at room temperature or overnight at 4° C with monoclonal Collagen I antibody (mAb39, 1:200 dilution) or monoclonal Laminin antibody (mAb52, 1:100 dilution) (Sarras et al., 1993). After primary antibody removal, samples were washed in PBS with 0.5% Tween 20 and 1% w/v BSA 3x for 10 minutes each. Secondary antibody staining was performed for 1 h at room temperature in blocking solution in the dark. Secondary antibodies used were Goat anti-Mouse Alexa Fluor 568 (Invitrogen A-11004) and 647 (Invitrogen A-21235). Secondary antibody was removed with three PBS washes. The second wash included Hoechst at 1:1000 dilution for nuclear staining (Invitrogen H3570). All steps were performed with gentle rocking. Samples were processed through 25%, 50%, and 80% glycerol/PBS washes, then placed between two coverslips (Fisher 12-541-025) separated by 0.12 mm spacer wells (Electron Microscopy Sciences 703279S), and both sides of each animal were imaged. Confocal images were obtained with Zeiss 980 Confocal with Airyscan 2 (see Image acquisition section).

Cryosection immunofluorescence: after warming slides, immunofluorescence was performed as with whole mount animals, but using primary antibody at 1:1000 dilution and no rocking. A hydrophobic barrier was drawn around the sections using a Liquid Blocker Super Pap Pen (Newcomer Supply 6505). Solutions were pipetted directly onto each slide, and incubations were carried out in a humid chamber. After staining, cryosectioned samples were placed under cover glass and sealed using nail polish.

### Marimastat and Dipyridyl treatment

Marimastat (Sigma-Aldrich M2699) was resuspended in DMSO at 50 mM and stored in small aliquots at −20° C to limit freeze/thaw cycles. Stock solutions were diluted 1:1000 in *Hydra* medium for a final concentration of 50 µM. 0.1% DMSO in *Hydra* medium was used as vehicle control. Dipyridyl (Sigma-Aldrich D216305) was resuspended in 100% ethanol at 100 mM, stored at −20° C, and diluted 1:1000 in *Hydra* medium for a final concentration of 100 µM. 0.1% ethanol in *Hydra* medium was used as vehicle control. During treatment, 16 animals per condition were kept in a single well of a six-well plate in 3 mL of media in the dark at 18° C. Animals were incubated in treatment or vehicle media for 1 h prior to injury for all experiments. For experiments lasting over 24 h, media was refreshed daily.

After fixation and PBS washes, animals were treated with 0.5% Triton-X in PBS for 15 minutes, then stained with Hoecht 33342 and AlexaFluor 594 Phalloidin (Invitrogen A12381) to aid in identification of small tentacle buds. Animals were mounted between two coverslips (as described in antibody staining section), and both sides of each animal were imaged to ensure complete quantification of tentacle buds.

### RNA probe synthesis

Primers for *collagen I* and *mmp* family members were designed using Primer3 (Untergasser et al., 2012) (Table S3). DNA fragments of 800-1200 base pairs in length were amplified from *Hydra vulgaris* cDNA using Platinum Taq polymerase and ligated into pGEM-T (Promega A3600) or pGEM-T Easy (Promega A1360) vectors. DIG or FITC-labeled probes were generated with DIG RNA Labeling Mix (Roche 11277073910).

### RNA fluorescent *in situ* hybridization (FISH)

FISH was performed following a previously published protocol (Little et al., 2025). Fixed animals were washed in PBS with 0.1% Tween 20 (PBT), then processed through a methanol gradient (33%, 66%, 100%) and stored in 100% methanol at −20° C overnight.

Samples were then rehydrated in a decreasing methanol (66%, 33%, 0%) gradient in PBT, permeabilized in 10 µg/ml proteinase K in PBT for 5 minutes, quenched in 4 mg/ml glycine, and washed three times in PBT for 5 minutes each. Samples were then washed in 0.1 M triethanolamine with increasing concentrations of acetic anhydride (0, 3, 6 µl/ml), and refixed in 4% PFA (3:1 ratio of PBT to 16% PFA) for 1 h at room temperature. After fixation, samples were washed in PBT and equilibrated in 2x SSC, then prehybridized in a mix of 2x SSC and hybridization solution (HS; 50% formamide, 5x SSC, 1x Denhardt’s, 100 μg/ml heparin, 0.1% Tween 20, 0.1% CHAPS) and moved to 56° C incubation during prehybridization. Samples were blocked for 2 h at 56° C in HS with 10 µl/ml sheared salmon sperm, then incubated with 180 ng of DIG or FITC-labeled RNA probes in 450 µl HS. Probes were denatured at 85° C for 5-10 minutes before adding to HS. Samples were incubated for approximately 60 h with gentle mixing approximately once per 24 h.

Probes were removed and samples washed at incubation temperature in decreasing HS concentrations in SSC (100% HS, 75%, 50%, 25%). They were then washed twice in 2x SSC with 0.1% CHAPS and moved to room temperature during the second wash, then washed four times in MABT (100 mM maleic acid, 150 mM NaCl, 0.1% Tween 20, pH 7.5) for 10 minutes each. Samples were blocked for 1 h at room temperature in MABT with 1% BSA, then for 2 h at 4° C in blocking buffer (80% MABT with 1% BSA / 20% sheep serum). Samples were incubated in anti-FITC-POD (Sigma-Aldrich 11426346910) or anti-DIG-POD (Sigma-Aldrich 11207733910) at 1:2000 dilution in blocking buffer at 4° C overnight.

Antibody was removed and samples were moved to room temperature and washed twice in MABT-BSA, then five times in MABT for 20 minutes per wash. Samples were then washed twice in 100 mM borate buffer with 0.1% Tween 20 for 5 minutes per wash, then stained with 75 μl tyramide solution (100 mM borate buffer, 2% dextran sulfate, 0.1% Tween 20, 0.003% H_2_O_2_, 0.15 mg/ml 4-iodophenol, 1:100 Alexa Fluor Tyramide 488 [Invitrogen B40953] or 594 [Invitrogen B40957]) for 25 min. Reaction was stopped with four PBT washes, then quenched in 100 mM glycine (pH 2.0) for 10 minutes and washed five more times for five minutes each in PBT. Samples were then incubated 10 minutes in Hoechst diluted 1:1000, processed through a glycerol gradient, and mounted in 80% glycerol on Fisher Brand Superfrost Plus slides using 0.12 mm depth spacer wells and localization was hand-counted by imaging with Leica DMIL LED Inverted Microscope. Confocal images were obtained with Zeiss 980 Confocal with Airyscan 2.

### Image acquisition

Confocal imaging was performed on a Zeiss 980 Confocal with Airyscan 2. For quantification of Collagen I staining and whole mount imaging of *Hydra*, slides were imaged using a 10x objective (Plan-Apochromat 10x/0.45). For quantification of Tg(*nanos*:eGFP)^tb1-in^ cells in endoderm, slides were imaged using a 20x objective (Plan-Apochromat 20x/0.8). Imaging parameters were set using ZEN software using two tracks, one to image Hoechst in the blue channel and one to image green and/or red channels. Images were taken at 1024×1024 pixel resolution with a pinhole size of 1 AU for imaging analysis. Z slice thickness was set to optimized thickness suggested by ZEN software based on Nyquist theorem. For occasional images for high resolution display, images were taken at 2048×2048 pixel resolution. For cryosections, a z stack was taken through each image to capture the thickness of the entire section. For whole mount animals, a z stack was taken from the ectodermal tissue in contact with the cover glass to the midway point of the animal’s thickness in the z plane, then the double coverslip mounting was flipped to image the opposite side of the animal.

For quantification of tentacle number in regeneration assays, animals were imaged on a Leica DMIL LED Inverted Microscope using the Hi Plan 4x/0.1 objective and EL6000 fluorescent light source with TXR ET filter cube. Images were captured using a Leica DFC300 G camera and LAS v4.7 software. Brightfield images of animals were acquired using a Leica M165 C stereo microscope with Leica MC170 HD (camera) using LAS v4.7 software.

### Image processing and quantification of GFP-positive cell invasion

For each section, a three channel (blue - nuclei, green - Tg(*nanos*:eGFP)^tb1-in^, red – Collagen I) three-dimensional stack was analyzed in FIJI (Schindelin et al., 2012). The endodermal portion of the regenerating head (defined as within 200 µm from the regenerating tip) was cropped using the Collagen I channel as a guide to distinguish epithelial layers. The endoderm of the Hoechst-stained channel was then fed into a custom watershed segmentation pipeline in CellProfiler to calculate total nuclei number (Carpenter et al., 2006). To obtain Tg(*nanos*:eGFP)^tb1-in^-positive cell counts, cells with fluorescence above background were marked by hand after randomizing file names to reduce bias. Cells were counted in two sections per animal. Endodermal eGFP-positive cell ratios were calculated as a simple ratio of eGFP-positive cells to total nuclei. Images are displayed in Figure 5 as maximum intensity projections.

### Image processing and quantification of Collagen I intensity and gap size

Single- channel (red – Collagen I) image stacks were analyzed in FIJI after generating maximum intensity projections. For cryosections, a segmented line 3 pixels wide was drawn along the ECM-depleted region in the regenerating head, then spline fitting was applied, and fluorescence intensity of the region was measured. Two similar lines of at least 200 µm each in length were drawn along each side of the uninjured region of the body column, and fluorescence intensity in the head region relative to the body was calculated. Three cryosections per animal were used to calculate average intensity in the head relative to body column for an individual animal, and eight animals per time point were used to calculate the time point average. For whole mount images, the Collagen I gap size was defined as the region of reduced staining intensity and irregular fibril orientation immediately oral to a line of saturated pixels on both imaged sides of each animal. This area was measured in FIJI and divided by the length across the body column at the base of the gap to obtain the average length from oral tip to base. Collagen I intensity in whole mounts was calculated by taking the average pixel intensity in the measured gap region and dividing by intensity in the uninjured portion of the animal body column.

### Transcript expression plots

Gene sequences were searched using BLAST against the *Hydra vulgaris* Jussy (Regeneration/Homeostatic spatial and loss of i-cells) database on HydrATLAS (https://hydratlas.unige.ch/blast/blast_link.cgi). After confirming sequence match, plots were downloaded and edited in Affinity Designer 2 to remove time course data not relevant to this study (Wenger et al., 2019).

### MMP domain analysis

We identified conserved domains in the *Hydra* MMP candidate sequences by sequence similarity to the National Center for Biotechnology Information (NCBI) Conserved Domain Database using Batch-CD Search (https://www.ncbi.nlm.nih.gov/Structure/bwrpsb/bwrpsb.cgi) with default settings. The identity and position of specific hits were recorded and annotated to create Figure S2. For some domains in sequences HVAEP1-G018597 and HVAEP1-G013033, domains were identified at the superfamily level, rather than specific hits; these domains are colored with a lighter shade in Figure S2. The identified domain for each *Hydra* MMP is given in Table S1. Putative propeptide domains were identified by manual search of the Hydra MMP amino acid sequences via the pattern PRCGXPD, where X is any amino acid. While the propeptide domain was recognizable in almost all MMP sequences, cnidarian sequences often showed minor divergence from the canonical bilaterian PRCGXPD pattern. The sequence of these putative cnidarian propeptide domains are given in Table S2.

### Statistical analysis

All experiments were performed with three biological replicates. Statistical significance was calculated by comparing independent populations with student’s non-paired t test.

## Acknowledgements

We thank Dr. Ming Zhang for his generous gift of Collagen I (mAb39) antibody. We also thank Dr. Bruce Draper and the UC Davis Light Microscopy Core for use of equipment.

