## Supplemental Figures for "Extracellular matrix remodeling supports *Hydra vulgaris* head regeneration and stem cell invasion"

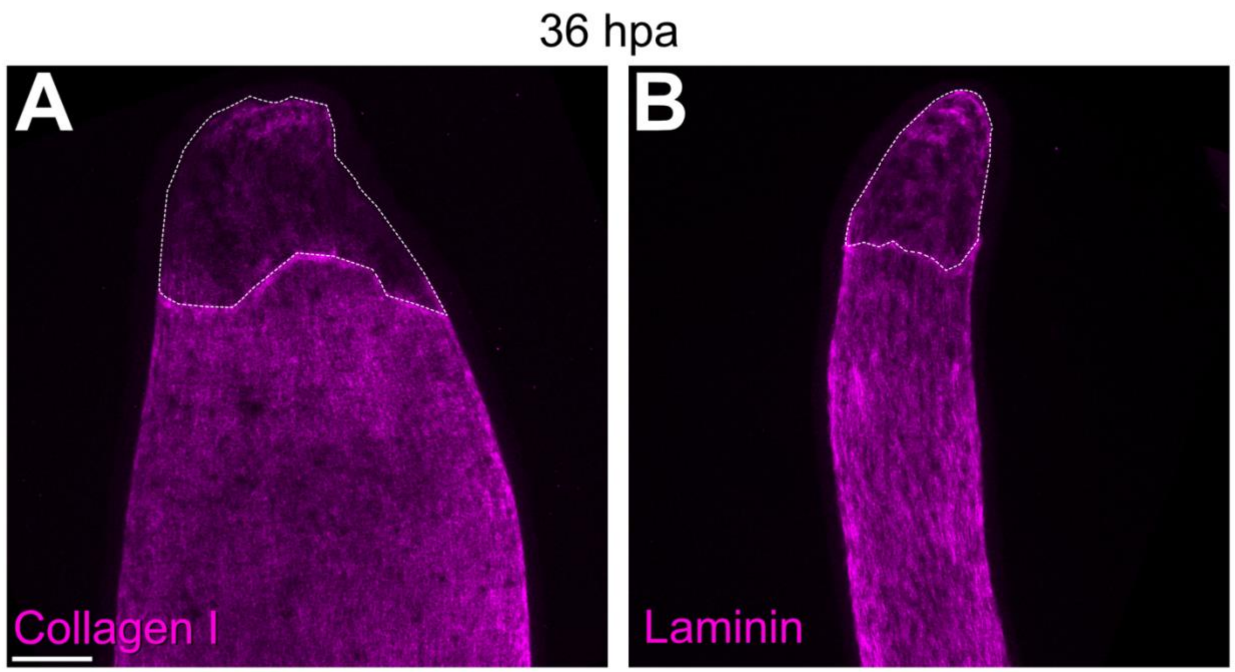


**Figure S1. Reduced levels of Collagen I and Laminin proteins in the regenerating *Hydra* head.** (A-B) Representative images of Collagen I (A) and Laminin (B) antibody staining in *Hydra* head regeneration at 36 hpa. Scale bar = 100 µm. White dashed line indicates the ECM protein-depleted region.

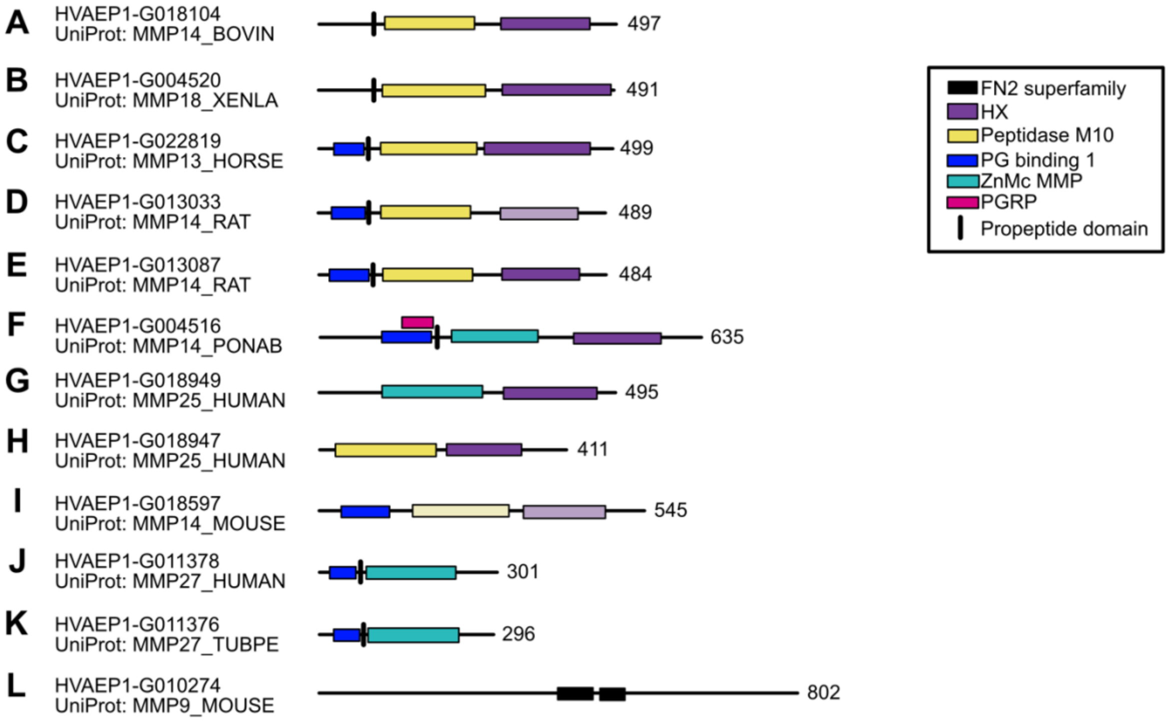


**Figure S2. Domain analysis of 12 transcripts with UniProt BLAST hits to MMP family members.** Previously published transcriptome data from our lab identified 12 transcripts whose best UniProt BLAST hit was an MMP from another species (Cazet et al., 2023; Siebert et al., 2019). We performed a conserved domain search to generate domain diagrams of each gene and identify the propeptide, catalytic (Zinc MMP or Peptidase M10), and hemopexin-like (HX) domains characteristic of MMP family members. We also noted domains present in some MMP family members (peptidoglycan [PG]-binding domain 1) or involved in peptidoglycan interactions (peptidoglycan recognition protein [PGRP] domain). (A-F) Six proteins contained an N-terminal propeptide, hemopexin-like domain, and a catalytic domain. (G-I) Three proteins lacked an N-terminal propeptide but contained the other two domains. (J-K) Two proteins lacked a hemopexin-like domain, but contained the other two domains. (L) G102074 did not include any of these domains but only fibronectin type II-like (FN2) domains, and we did not analyze it further. Lighter shaded domains in G013033 (D) and G018597 (I) indicate that the domain was classified at the superfamily level, which indicates a broad relationship to a conserved domain superfamily but the absence of a high-confidence hit to any specific conserved domain model within the superfamily. At the left of each domain map is the gene number and UniProt BLAST hit; at right, the number of amino acid residues.

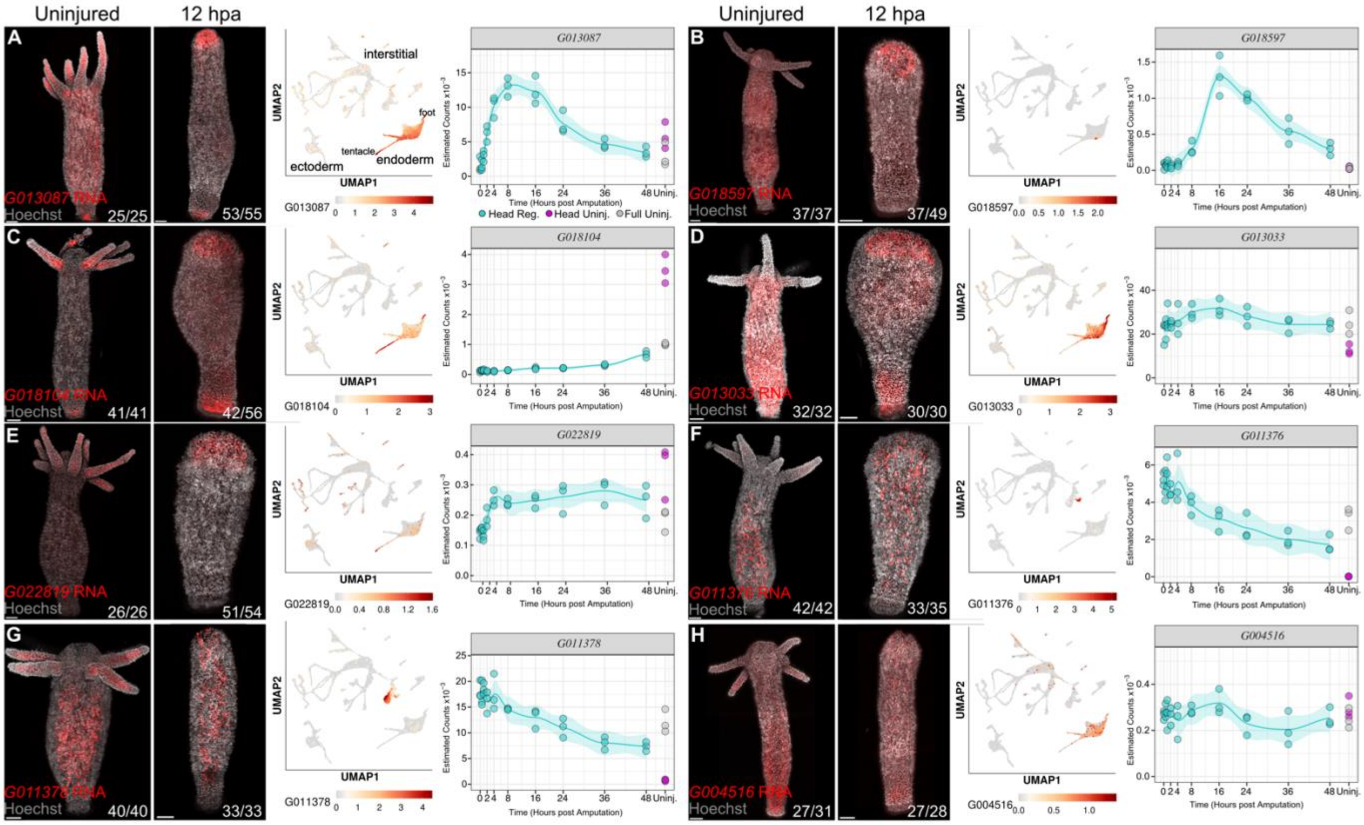


**Figure S3. A subset of *mmp* transcripts are expressed in the regenerating *Hydra* head by FISH.** (A-E) Five separate *mmp* transcripts lack head-specific expression in the uninjured animal (first panel) but localize to the regenerating head endoderm at 12 hpa (second panel). Previously published scRNA-seq maps (third panel) of uninjured animals show agreement between these data and FISH results (Cazet et al., 2023; Siebert et al., 2019). Endoderm-specific expression in regenerating heads of *G013087* and *G018597* transcripts (A-B) is accompanied by a significant increase in transcripts in a bulk RNA-seq data set (fourth panel) (Wenger et al., 2019). *G018014, G013033*, and *G022819* transcripts show more modest increases in overall transcript counts later in the regeneration time course (C-E). The *mmp* transcripts that do not show head-specific localization during regeneration are primarily expressed in interstitial lineage cells, and *G011376* and *G011378* show declines in transcript counts during regeneration (F-H). All scale bars = 100 µm. Ratio in bottom right hand corner of each FISH image indicates number of animals showing depicted expression out of total stained animals. Panel A is repeated from Figure 3 to show all *mmp* transcript probes in a single figure.

### Supplemental Table Legends

Table available as separate file.

**Table S1. Full Hit Data Table Produced by a Conserved Domain Search of 12 Candidate Hydra MMPs.** The full hit data output table of an NCBI conserved domain search for the list of 12 Hydra MMP candidates. Query sequence identity (column A), hit type (column B), position (columns D, E) and accession (column H) of each domain hit is given. A full description of the data described in each column can be found at https://www.ncbi.nlm.nih.gov/Structure/cdd/cdd_help.shtml#BatchRPSBDownloadDomainHits.

| Sequence | Propeptide Sequence | Position |
|---|---|---|
| **HVAEP1-G010274** | NA | NA |
| **HVAEP1-G022819** | PRCGIED | 84-90 |
| **HVAEP1-G018597** | NA | NA |
| **HVAEP1-G004516** | PRCGVSD | 200-206 |
| **HVAEP1-G004520** | PRCGMPD | 92-98 |
| **HVAEP1-G018104** | PRCGLPD | 92-98 |
| **HVAEP1-G013033** | PRCGLKD | 81-87 |
| **HVAEP1-G013087** | PRCGLPD | 86-92 |
| **HVAEP1-G018947** | NA | NA |
| **HVAEP1-G018949** | NA | NA |
| **HVAEP1-G011378** | PRCGISD | 72-78 |
| **HVAEP1-G011376** | PRCGVSD | 72-78 |

**Table S2. Propeptide sequences identified in *Hydra* MMP candidates.** This table shows the gene number of each *Hydra* MMP candidate (column A), propeptide sequence PRCGXPD (where X is any amino acid) or closest hit (column B) and its location within the amino acid sequence (column C). G004520, G018104, and G013087 contain the full PRCGXPD sequence, while G022819, G004516, G013033, G011378, and G011376 have an additional amino acid substitution at the sixth position. A propeptide domain was not found in G010274, G018597, G018947, or G018949.

| Gene ID | UniProt BLAST Hit | Forward | Reverse |
|---|---|---|---|
| G022819 | MMP13_HORSE | ACAGGGTGGAAAATGGCATA | taatacgactcactatagggACGTTAGTGGGTAGCCATCG |
| G018597 | MMP14_MOUSE | GAACCGGGTAAATACGCAGA | atttaggtgacactatagCCAATTGCAGATGAAGGTGA |
| G004516 | MMP14_PONAB | GGTTTGTGGATGGCAGAAAT | atttaggtgacactatagTGCATCCACATCTCGAAAAC |
| G018104 | MMP14_BOVIN | CGAGTTGGACAACTGCTTGA | taatacgactcactatagggGATGGTACATTGCCCCAGAA |
| G013033 | MMP14_RAT | GCACATGCGTTTTACCCTTT | atttaggtgacactatagGCACGTTCTTTGGAGCTTTC |
| G013087 | MMP14_RAT | GACGACCCCTACAGGTTTGA | taatacgactcactatagggCGGAAAGTTCAACCAAGCTC |
| G011378 | MMP27_HUMAN | TTGGGCTTTTTGACTCTATGTG | atttaggtgacactatagTATGCACAACCGTACGCAAT |
| G011376 | MMP27_TUPBE | GTATTTGTATGGACAATCCGACGA | taatacgactcactatagggCTTTCCAATGATCCCAAGTACA |
| G020291 | CO2A1_CHICK | AGGATCTCGAGGTGCTGATG | taatacgactcactatagggTGTGGTCCTTGTGATCCTCC |

**Table S3. Primer sequences used for FISH probe template synthesis.** This table shows the forward and reverse primers used for amplification of 800-1200 base pair probe templates from *Hydra* cDNA. Column A indicates the Gene ID, and Column B the gene name. Columns C and D contain the primer sequences. Capital letters indicate the binding sequences, while lowercase letters in Column D represent either the SP6 or T7 polymerase binding sequence added to the PCR product.
